# Evidence for a Spoken Word Lexicon in the Auditory Ventral Stream

**DOI:** 10.1101/2022.10.09.511436

**Authors:** Srikanth R. Damera, Lillian Chang, Plamen P. Nikolov, James A. Mattei, Suneel Banerjee, Laurie S. Glezer, Patrick H. Cox, Xiong Jiang, Josef P. Rauschecker, Maximilian Riesenhuber

**Author notes:** Correspondence should be addressed to M.R. Department of Neuroscience, Georgetown University Medical Center, Research Building Room WP-12, 3970 Reservoir Rd. NW, Washington, DC 20007, USA.

## Abstract

The existence of a neural representation for whole words (i.e., a lexicon) is a common feature of many models of speech processing. Prior studies have provided evidence for a visual lexicon containing representations of whole written words in an area of the ventral visual stream known as the “Visual Word Form Area” (VWFA). Similar experimental support for an auditory lexicon containing representations of spoken words has yet to be shown. Using fMRI rapid adaptation techniques, we provide evidence for an auditory lexicon in the “Auditory Word Form Area” (AWFA) in the human left anterior superior temporal gyrus that contains representations highly selective for individual spoken words. Furthermore, we show that familiarization with novel auditory words sharpens the selectivity of their representations in the AWFA. These findings reveal strong parallels in how the brain represents written and spoken words, showing convergent processing strategies across modalities in the visual and auditory ventral streams.

**Highlights:**

- Individual auditory word form areas (AWFA) were defined via an auditory localizer
- The AWFA shows tuning for individual real words but not untrained pseudowords
- The AWFA develops tuning for individual pseudowords after training

## Introduction

Speech perception is perhaps the most remarkable achievement of the human auditory system and one that likely is critically dependent on its overall cortical architecture. It is generally accepted that the functional architecture of auditory cortex in human and nonhuman primates comprises two processing streams^1–3^. There is an auditory dorsal stream that is involved in the processing of auditory space and motion^4^ as well as in sensorimotor transformations such as those required for speech production^5–8^. There is also an auditory ventral stream specialized for recognizing auditory objects such as spoken words. This stream is organized along a simple-to-complex feature hierarchy^2^, akin to the organization of the visual ventral stream^9^.

Visual object recognition studies support a simple-to-complex model of cortical visual processing in which neuronal populations in the visual ventral stream are selective for increasingly complex features and ultimately visual objects along a posterior-to-anterior gradient extending from lower-to-higher-order visual areas^10,11^. For the special case of recognizing written words, this simple-to-complex model predicts that progressively more anterior neuronal populations are selective for increasingly complex orthographic patterns ^12,13^. Thus, analogous to general visual processing, orthographic word representations are predicted to culminate in representations of whole visual words – an orthographic “lexicon”. Evidence suggests that these lexical representations are subsequently linked to concept representations in downstream areas like the anterior temporal lobe ^14–17^. The existence of this orthographic lexicon in the brain is predicted by neuropsychological studies of reading ^18^. Indeed, functional magnetic resonance imaging (fMRI) ^19,20^ and, more recently, electrocorticographic (ECoG) data ^21,22^ have confirmed the existence of such a lexicon in a region of the posterior fusiform cortex known as the Visual Word Form Area (VWFA) ^12,23^.

It has been proposed that an analogous simple-to-complex hierarchy exists in the auditory ventral stream as well^24,25^ extending anteriorly from Heschl’s Gyrus along the superior temporal cortex (STC)^2,26^. Yet, the existence and location of a presumed “auditory lexicon”, *i*.*e*., a neural representation for the recognition (and storage) of real words, has been quite controversial ^27^: The traditional, “posterior” view is that the auditory lexicon should be found in posterior superior temporal cortex (pSTC)^28^. In contrast, a notable meta-analysis ^26^ provided strong evidence for the existence of word-selective auditory representations in *anterior* STC (aSTC), consistent with imaging studies of speech intelligibility ^29,30^ and proposals for an “auditory word form area” (AWFA) in the human left anterior temporal cortex^26,31^. Such a role of the aSTC is compatible with nonhuman primate studies that show selectivity for complex communication calls in aSTC ^32–34^ and demonstrate, in humans and nonhuman primates, that progressively anterior neuron populations in the STC pool over longer timescales ^35–38^. In this “anterior” account of lexical processing, the pSTC and speech-responsive regions in the IPL are posited to be involved in “inner speech” and phonological reading (covert articulation), but not auditory comprehension^6,39^. Yet, despite this compelling alternative to traditional theories, there is still little direct evidence for an auditory lexicon in the aSTC.

Investigating the existence and location of auditory lexica is critical for understanding the neural bases of speech processing and, consequently, the neural underpinnings of speech processing disorders. However, finely probing the selectivity of neural representations in the human brain with fMRI is challenging, in part because it is difficult to assess the selectivity of these populations. Many studies have identified speech processing areas by contrasting speech stimuli with various non-speech controls ^29,40,41^. However, these coarse contrasts cannot reveal what neurons in a particular auditory word-responsive ROI are selective for, *e*.*g*., phonemes, syllables, or whole words. More sensitive techniques such as fMRI rapid adaptation (fMRI-RA ^42,43^) are needed to probe the selectivity of speech representations in the brain and resolve the question of the existence of auditory lexica. In the current study, we used fMRI-RA to test the existence of lexical representations in the auditory ventral stream. Paralleling previous work in the visual system that used fMRI-RA to provide evidence for the existence of an orthographic lexicon in the VWFA ^19,20,44^, we first performed an independent auditory localizer scan that we used to identify the AWFA in individual subjects, and then conducted three fMRI-RA scans that probed the representation in the AWFA and its plasticity. The first two scans consisted of real words and pseudowords (pronounceable nonwords), respectively. These scans revealed an adaptation profile consistent with lexical selectivity in the putative AWFA for real words, but not novel pseudowords, directly replicating results for written words in the VWFA. We then tested the lexicon hypothesis by predicting that training subjects to recognize novel pseudowords would add them to their auditory lexica, leading them to exhibit lexical-like selectivity in the AWFA following training, as previously shown for written words in the VWFA ^19^. To do so, we conducted a third fMRI-RA scan after pseudoword training. Results from this scan showed real word-like lexical selectivity to the now-familiar pseudowords following training, supporting the role of the AWFA as an auditory lexicon shaped by experience with auditory words.

## Results

### Auditory Localizer Scan Identifies Bilateral Speech-Selective ROI in the Auditory Ventral Stream

An independent auditory localizer scan was used to identify a putative auditory word form area (AWFA) near previous literature coordinates (Fig. 2A). To do so, analogous to prior studies of written word representations in the VWFA ^19,20,44^, we first identified group-level auditory speech-selective areas (see Materials and Methods) by examining the “(*Real Words) vs*.*Silence”* contrast thresholded at p≤0.001 masked by the “*RW vs. Scrambled Real Words”* contrast thresholded at p≤0.05 at the group-level cluster-corrected at the FDR p ≤ 0.05. This revealed several clusters of activation in the superior temporal, frontal, and inferior temporal cortices (Fig. 2B). A local peak in the left STG was identified at MNI: -62, - 14, 2 (Fig. 2C) near the literature coordinates^26^ of MNI: -61, -15, -5. Individual subject AWFA ROIs (Fig. 2D) were created by building a 50-voxel ROI around the local peak of each subject (mean ± std: -62 ± 2, -14.9 ± 3, 2.6 ± 2.6) closest to the group peak (see Materials and Methods).

**Figure 1:**
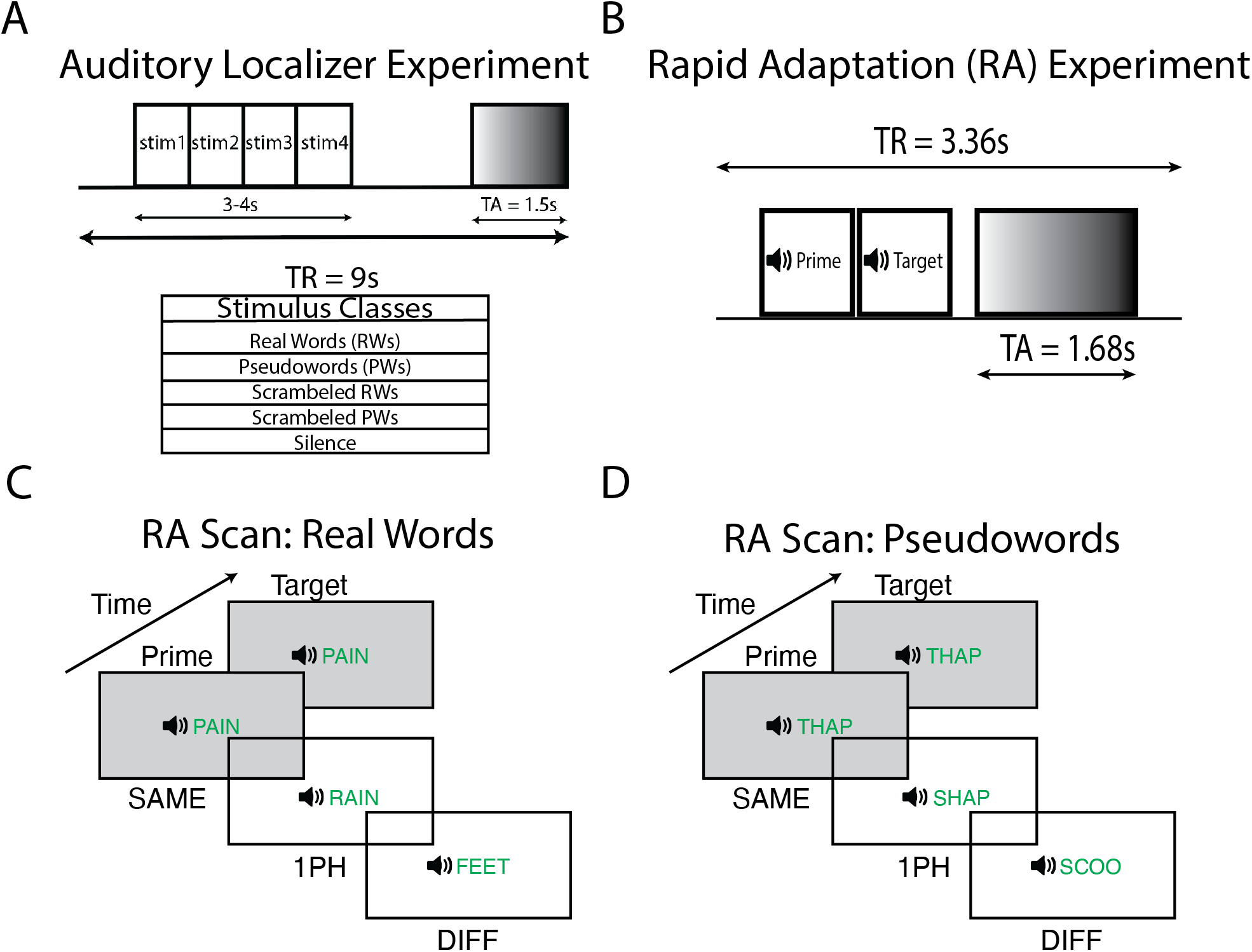
Rapid adaptation and Auditory Localizer Experimental Paradigms. (A) Shows the slow clustered acquisition paradigm used in the auditory localizer scan. Each trial was 9s long with 1.5s of volume acquisition and 7.5s of silence. During the silent period, the subject heard 4 sounds from one of 5 stimulus classes and performed a 1-back task. (B) Shows the rapid clustered acquisition paradigm used for the RA scans ^45,46^. Each trial was 3.36s long with 1.68s of volume acquisition. During the silent period, two spoken words were played to the subject with a 50ms ISI. The first word acted as a prime and the second word the target. (C and D) Shows the experimental paradigm for Real Words (RW) and Pseudowords (UTPW/TPW), respectively. The prime was followed by a target word that was either the same word (SAME), a word that differed from the target by one phoneme (1PH), or a word that shared no phonemes with the target (DIFF). Furthermore, subjects were presented with silence trials that served as an explicit baseline. During the task, subjects were asked to attend to all the words and respond when they heard the oddball stimulus (RW or PW containing the rime ‘-ox’ e.g., “socks”) in either the prime or target position.

**Figure 2:**
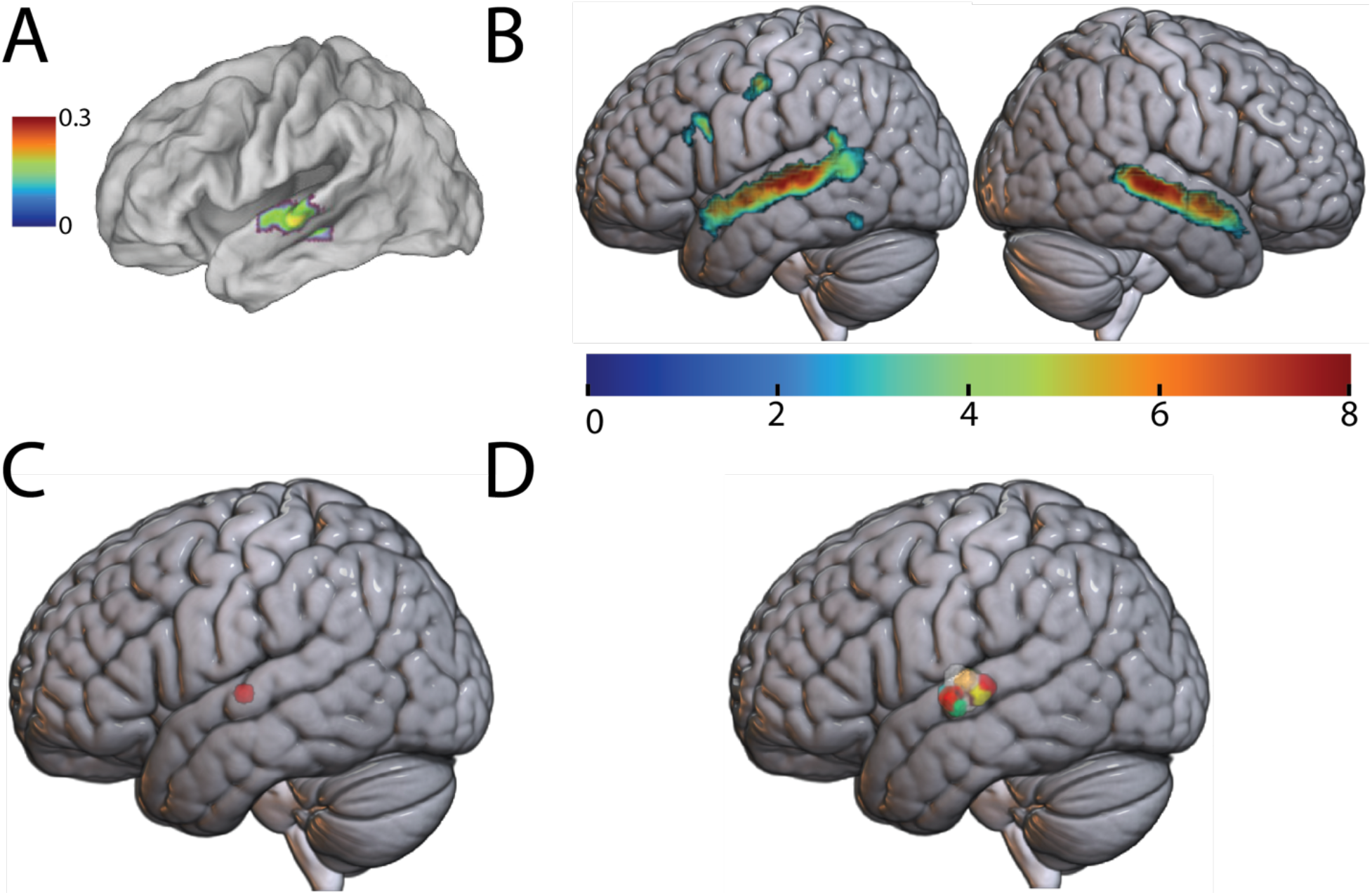
Identifying the Auditory Word Form Area (AWFA). (A) Proposed location of the Auditory Word Form Area (MNI: -61, -15, -5) adapted from ^26^. Color bar reflects the ALE ^47^ statistic. (B) Shows the “*RW-Silence*” contrast (p ≤ 0.001) masked by the “*RW-Scrambled Real Words”* contrast (p ≤ 0.05) in the auditory localizer scan. Only clusters significant at the FDR p ≤ 0.05 level are shown. Colors reflect t-statistics. (C) Marks the peak in the left STG (MNI: -62, -14, 2). (D) Shows the Auditory Word Form Area defined in individual subjects.

**Supplementary Table 1.** Individual Subject AWFA ROIs

| Subjects | X-Coordinate | Y-Coordinate | Z-Coordinate |
| --- | --- | --- | --- |
| sub-MR1076 | -62 | -16 | 2 |
| sub-MR1106 | -62 | -16 | 4 |
| sub-MR1215 | -62 | -16 | 4 |
| sub-MR1216 | -62 | -12 | 0 |
| sub-MR1217 | -64 | -20 | 4 |
| sub-MR1220 | -58 | -16 | 0 |
| sub-MR1221 | -64 | -14 | 4 |
| sub-MR1222 | -64 | -10 | 0 |
| sub-MR1233 | -62 | -12 | 8 |
| sub-MR1234 | -62 | -12 | 6 |
| sub-MR1240 | -66 | -16 | 6 |
| sub-MR1242 | -62 | -16 | 0 |
| sub-MR1252 | -60 | -10 | 2 |
| sub-MR1253 | -60 | -10 | 0 |
| sub-MR1258 | -58 | -12 | 0 |
| sub-MR1272 | -62 | -18 | 2 |
| sub-MR1273 | -58 | -18 | 6 |
| sub-MR1277 | -64 | -18 | 2 |
| sub-MR1278 | -66 | -22 | 4 |
| sub-MR1282 | -60 | -12 | -2 |
| sub-MR1283 | -62 | -18 | 4 |
| sub-MR1284 | -60 | -16 | 4 |
| sub-MR1286 | -64 | -12 | -2 |
| sub-MR1287 | -60 | -16 | 6 |
| sub-MR1289 | -66 | -20 | 2 |
| sub-MR1294 | -62 | -10 | 2 |

### Lexical Selectivity for Real Words but not Pseudowords in the AWFA

The first two fMRI-RA scans were performed with real words (RW) and pseudowords (“untrained pseudowords”, UTPW), respectively. In these experiments, we predicted the lowest signal for the “SAME” condition since the two identical words presented in that condition would repeatedly activate the same neural populations, thereby causing maximal adaptation. Likewise, we predicted the least amount of adaptation for the “DIFF” condition because two words that share no phonemes should activate disjoint groups of neurons, irrespective of whether responsiveness to auditory words in the localizer scan in that ROI was due to neurons selective for phonemes, syllables or whole words. Finally, we tested specific predictions regarding responses in the 1PH condition. Specifically, if neurons in the AWFA ROI are tightly tuned to whole real words (i.e., if the AWFA contains an auditory lexicon), then, as in the DIFF condition, the two similar but non-identical real words in the 1PH condition should have minimal neural overlap and therefore no adaptation should occur and response levels in the 1PH condition should be comparable to that of the DIFF condition (as found for written real words in the VWFA^20^). In contrast, if neurons in the AWFA were tuned to sublexical phoneme combinations, then there should be a gradual release from adaptation from SAME to 1PH to DIFF, with 1PH < DIFF, as sublexical overlap would continue to increase from 1PH to DIFF. For pseudowords, we predicted that there would be a gradual increase in the BOLD signal paralleling the increasing dissimilarity (i.e., SAME to 1PH to DIFF). This is thought to reflect low-level activation of RW-tuned neurons to phonologically similar pseudowords. These predictions mirror findings in the VWFA for written words^19,20,44^ and are compatible with an experience-driven increase in selectivity of neurons in the AWFA to real words because of extensive auditory experience with and the need to discriminate among real words but not pseudowords.

To test our hypotheses, we ran a 2-way repeated measures ANOVA on AWFA responses to investigate the relationship between lexicality (real words, RW, and untrained pseudowords, UTPW) and word similarity (SAME, 1PH, and DIFF). The ANOVA was run on subjects that successfully completed both the RW and UTPW scans (n=24). This revealed a significant main effect of similarity (*F*_2,46_ = 42.597; *p* = 3.4E-11) but no significant main effect of lexicality (*F*_1,23_ = 2.547; *p* = 0.124). Critically, however, the analysis revealed a significant interaction between lexicality and word similarity (*F*_2,46_ = 4.092; *p* = 0.023). Planned paired t-tests revealed an adaptation profile in the AWFA that was consistent with tight neural tuning for individual real words (Fig. 3A): There was a significant difference in the mean percent signal between the DIFF vs. SAME conditions (*t(23)* = 7.86; *p* <0.001) and the 1PH vs. SAME (*t(23)* = 5.71; *p* < 0.001). However, there was no significant difference between the DIFF and 1PH conditions (*t(23)* = 1.95; *p* = 0.189). Similar results were obtained using all 26 subjects who completed the RW scan (Supplementary Fig. 1). In contrast, the adaptation profile for UTPWs was not consistent with lexical selectivity (Fig. 3B): There was a significant response difference between the DIFF vs. SAME conditions (*t(23)* = 7.03; *p* < 0.001), the 1PH vs. SAME (*t(23)* = 3.28; *p* = 0.010), and, critically, also for the DIFF vs. 1PH conditions (*t(23)* = 3.89; *p* = 0.002). Finally, we ran a whole-brain analysis to test if lexical representations were localized to the anterior STG or more distributed in nature. Specifically, for RW the conjunction of DIFF vs. SAME and 1PH vs. SAME after excluding voxels where DIFF vs. PH was p < 0.05 produced a cluster in the left anterior STG (FWE-corrected p < 0.05; MNI: -62, -14, 6) within 3.5mm from the average coordinate (MNI: -62, -14.9, 2.6) of our individual ROIs. Importantly, this analysis for UTPWs produced no significant clusters. Thus, the whole-brain analysis also supports the special status of the AWFA as the location of a lexicon for spoken real words.

**Figure 3:**
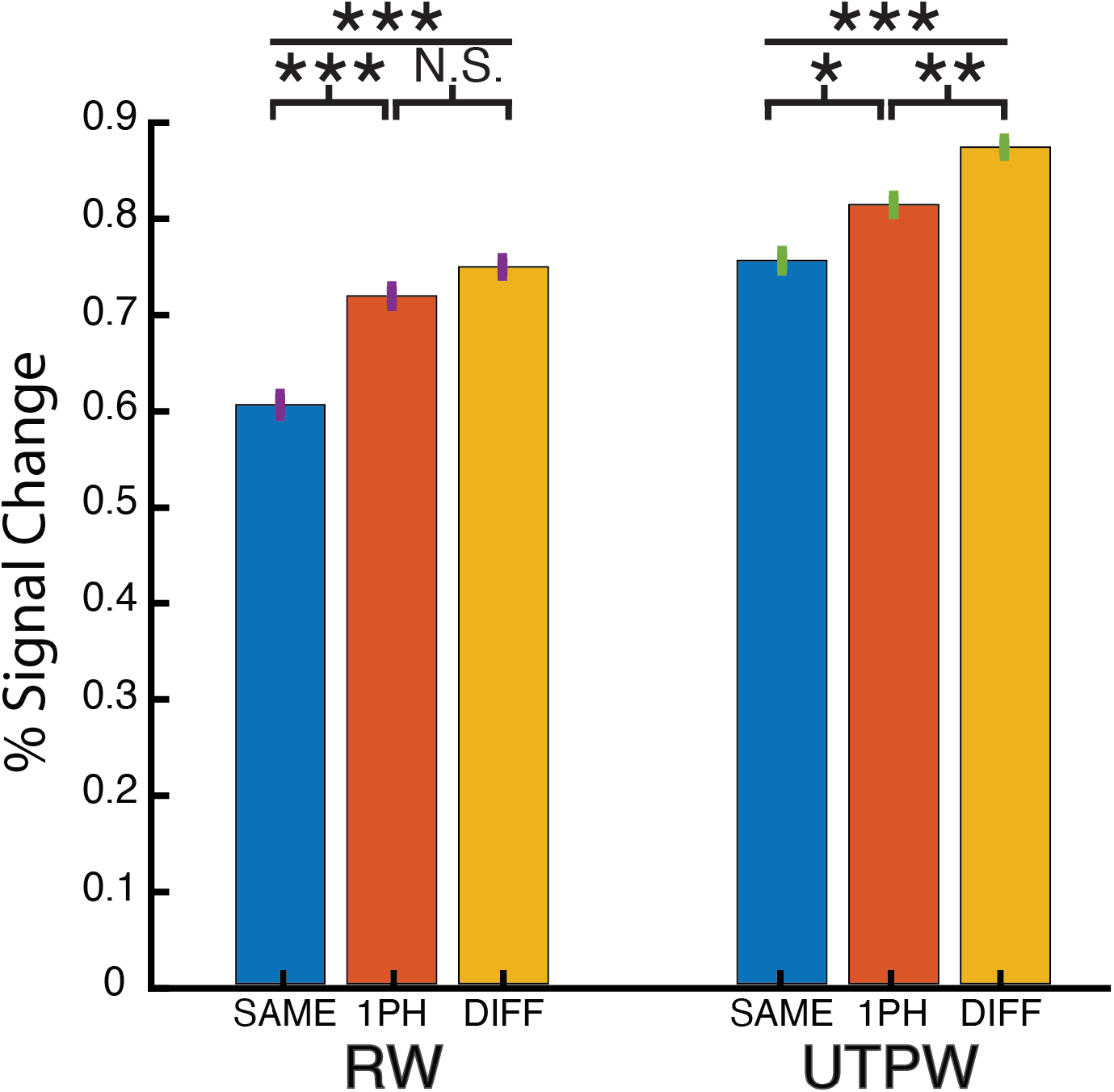
Evidence for Auditory Lexical Representations in the AWFA. Within-subject (n=24) adaptation profile for auditory real words (RWs) and untrained pseudowords (UTPWs). Patterns of release from adaptation are compatible with tight tuning to individual real words consistent with an auditory lexicon. In contrast, UTPWs show a graded release from adaptation as a function of phonological similarity. ***, **, *, and N.S. mark: *p* ≤0.001, ≤0.01, ≤0.05, and not significant (>0.1), all Bonferroni-corrected for multiple comparisons.

Planned paired t-tests revealed an adaptation profile that was consistent with tight neural tuning for individual RWs (SFig. 1). There was a significant difference in the mean percent signal between the DIFF vs. SAME conditions (*t(25)* = 6.94; p <0.001) and the 1PH vs. SAME (*t(25)* = 5.36; p < 0.001). However, there was no significant difference between the DIFF and 1PH conditions (*t(25)* = 1.70; p = 0.3039).

**Supplementary Figure 1:**
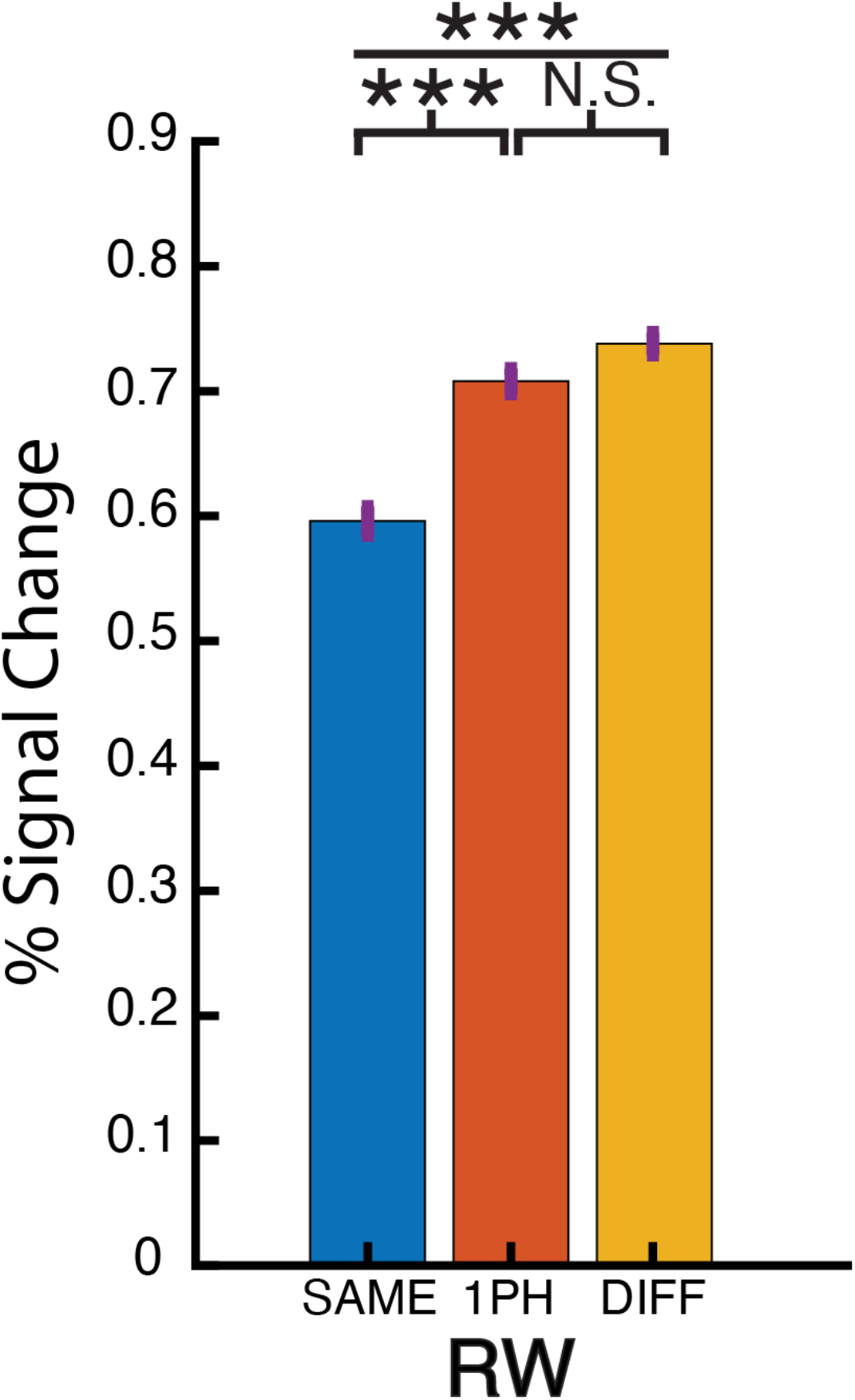
Evidence for Auditory Lexical Representations in the AWFA. Adaptation profile for auditory real words (RWs) using the full dataset (n=26). RWs show tight tuning to individual real words consistent with an auditory lexicon. ***, **, *, and N.S. mark: p ≤0.001, ≤0.01, ≤0.05, and not significant (>0.1), all Bonferroni-corrected for multiple comparisons.

### Adaptation Patterns to Pseudowords in the AWFA Exhibit Lexical Selectivity After, but not Before Familiarization

Next, in the pre- and post-training scans (UTPW and TPW, respectively), we tested the hypothesis that familiarization with previously novel pseudowords drives the formation of lexical selectivity in the AWFA. To do so, we examined the adaptation profiles for RWs, pseudowords before training (untrained pseudowords, UTPW), and pseudowords after training (trained pseudowords, TPW) in subjects who had completed all three scans (n = 16). We ran a 2-way repeated measures ANOVA to investigate the relationship between lexicality (RW, UTPW, and TPW) and similarity (SAME, 1PH, and DIFF). This revealed a significant main effect of similarity (*F*_2,30_ = 43.023; *p* = 2.95E-9) but not a significant main effect of lexicality (*F*_2,30_ = 2.398; *p* = 0.116). Critically, there was again a significant interaction between lexicality and similarity (*F*_4,60_ = 4.144; *p* = 0.012).

Consistent with the full dataset (Fig. 3), planned paired t-tests revealed an adaptation profile that was consistent with tight neural tuning for individual RWs (Fig. 4). There was a significant difference in the mean percent signal change between the DIFF vs. SAME conditions (*t(15)* = 7.28; *p* < 0.001) and 1PH vs. SAME (*t(15)* = 5.85; *p* < 0.001). However, there was no significant difference between the DIFF and 1PH conditions (*t(15)* = 0.750; *p* = 1). In contrast, the adaptation profile for UTPW was not consistent with lexical selectivity (Fig. 4): There was a significant difference between the DIFF vs. SAME conditions (*t(15)* = 7.84; *p* < 0.001) and the DIFF vs. 1PH conditions (*t(15)* = 4.57; *p* = 0.001), but not 1PH vs. SAME (*t(15)* = 2.16; *p* = 0.141). Crucially, the adaptation profile for PW *after* training (i.e., TPW) was consistent with lexical selectivity (Fig. 4): There was a significant difference in the mean percent signal between the DIFF vs. SAME conditions (*t(15)* = 5.55; *p* < 0.001) and the 1PH vs. SAME (*t(15)* = 3.17; *p* = 0.019). However, there was no significant difference between the DIFF and 1PH conditions (*t(15)* = 1.71; *p* = 0.327).

**Figure 4:**
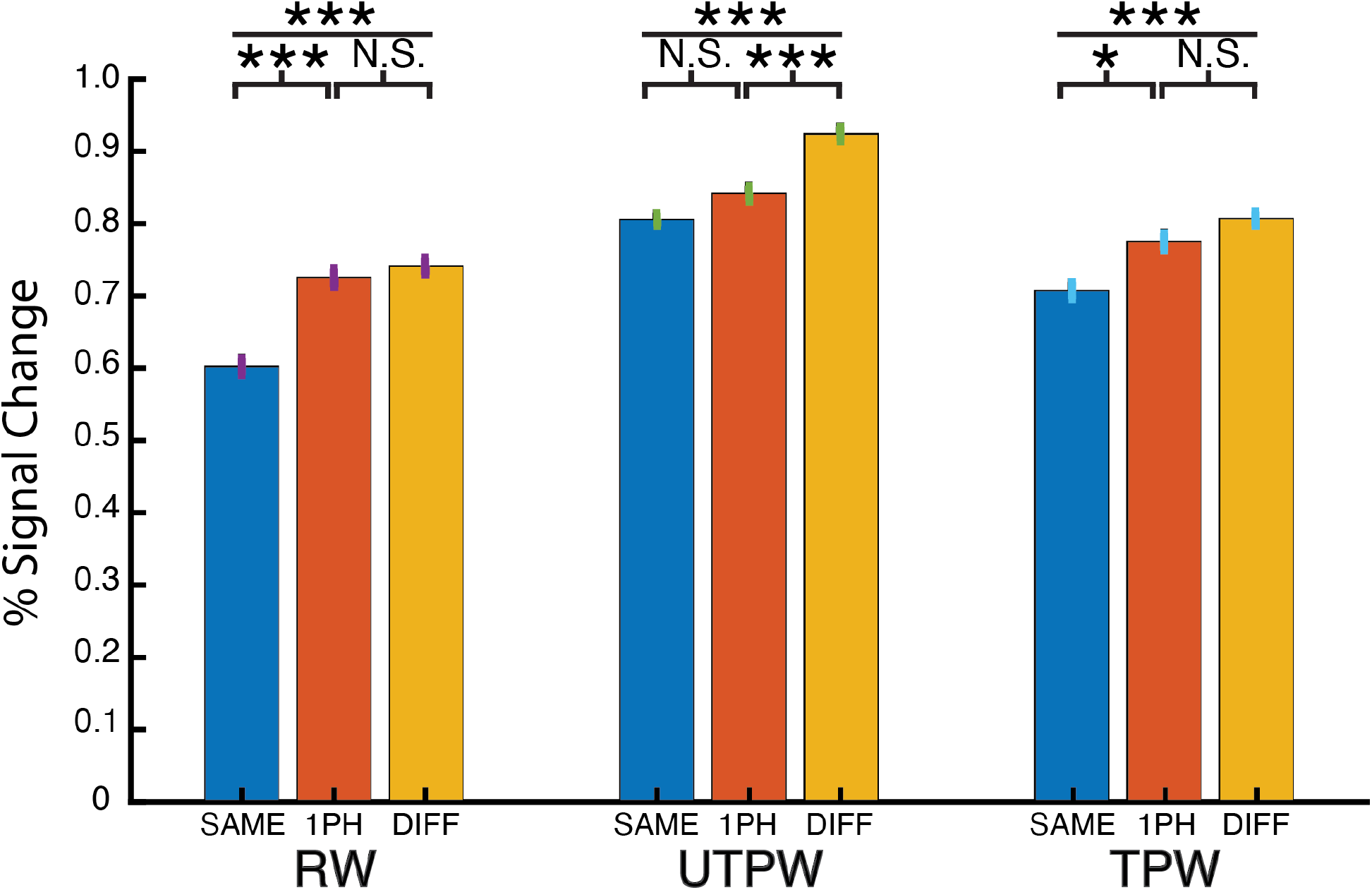
Auditory Lexical Representations Emerge in the AWFA for Pseudowords After Familiarization Training. Within-subject (n = 16) adaptation profile for auditory real words (RW), untrained pseudowords (UTPW), and trained pseudowords (TPW). RW adaptation profile shows tuning to individual real words consistent with an auditory lexicon. Untrained pseudowords (UTPW) show a graded release from adaptation as a function of phonological similarity. Importantly, following familiarization training, adaptation patterns in the AWFA to the same pseudowords (now trained, TPW) reveal tight lexical tuning, similar to RW. ***, **, *, and N.S. mark: *p* ≤0.001, ≤0.01, ≤0.05 and not significant (>0.1), all Bonferroni-corrected.

### The AWFA Connects to the Language Network

We next calculated task-based functional connectivity in the auditory localizer dataset (n=26) to examine the connections between the AWFA and the rest of the brain. This seed-to-voxel analysis (Fig. 5) showed that the AWFA is highly connected with brain regions previously shown ^48^ to be involved in language processing such as the inferior frontal gyrus (local peak at MNI -46, 12, 26) and premotor cortex (local peak at MNI -50, -8, 48). Especially noteworthy is the connectivity between the AWFA and a cluster in the left posterior fusiform cortex (circled in red; MNI: -44, -46, -14) that encompases the reported location of the VWFA (MNI: -45, -54, -20) ^12,49^.

**Figure 5:**
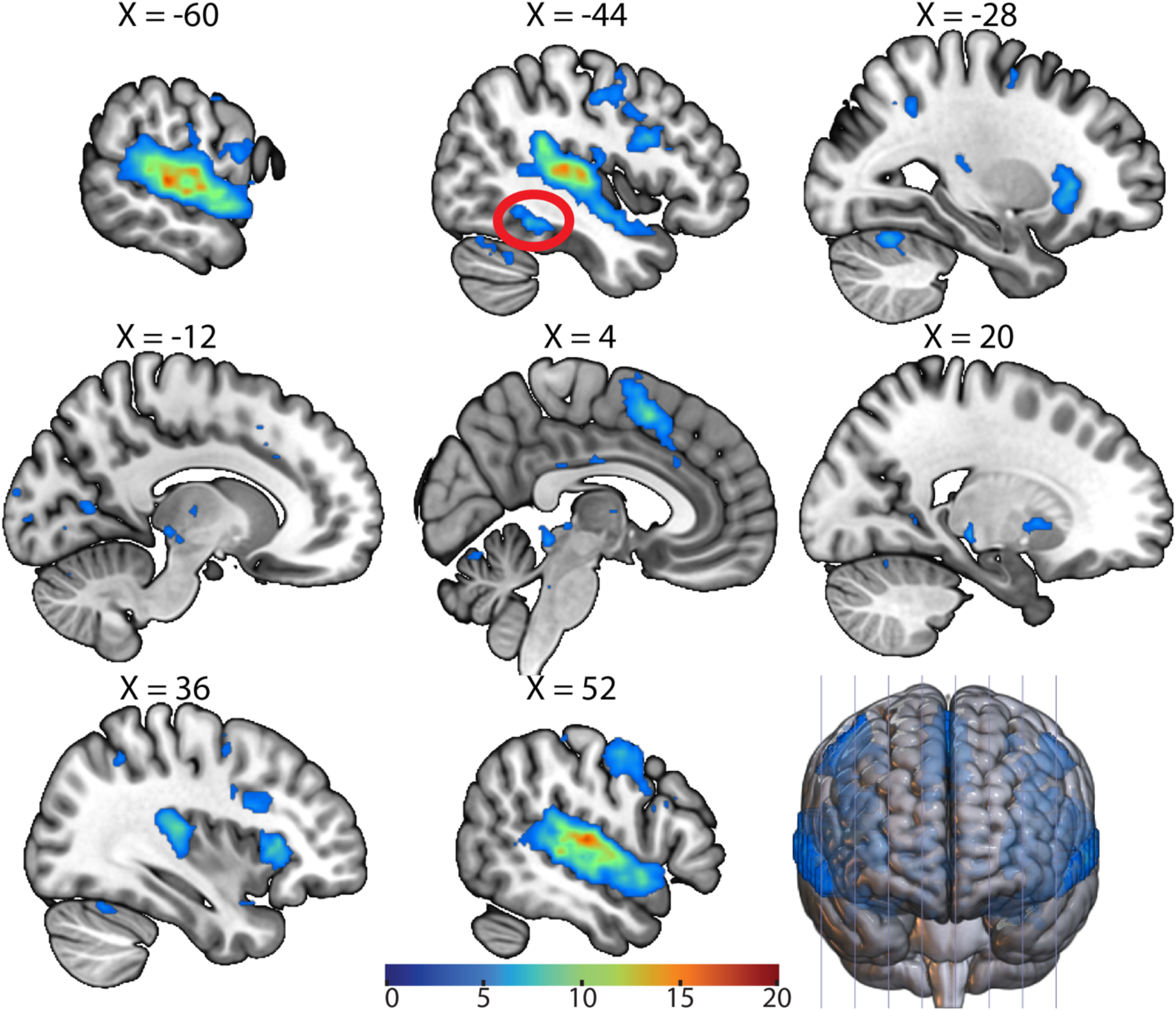
Functional Connectivity of the AWFA. Whole-brain functional connectivity of the AWFA during the auditory localizer task (n=26). Results are thresholded at a voxel-wise *p* ≤ 0.001 and cluster-level *p* ≤ 0.05, FWE corrected. Cluster corresponding to the literature coordinates of the VWFA is circled in red.

**Supplementary Table 2.** Significant Clusters of Connectivity With the AWFA

| X-Coordinate | Y-Coordinate | Z-Coordinate | size | p-FDR |
| --- | --- | --- | --- | --- |
| 64 | -20 | 0 | 5239 | 0 |
| -64 | -28 | 8 | 5217 | 0 |
| -6 | 12 | 48 | 970 | 0 |
| 50 | -2 | 50 | 934 | 0 |
| -12 | 26 | 58 | 800 | 0 |
| -46 | 12 | 26 | 799 | 0 |
| -40 | -76 | 36 | 737 | 0 |
| -34 | 24 | 8 | 429 | 0 |
| 18 | -24 | -4 | 351 | 0 |
| -26 | -62 | -24 | 272 | 0 |
| 46 | -68 | 28 | 258 | 0 |
| 12 | -70 | -18 | 256 | 0 |
| 34 | 30 | 54 | 242 | 0 |
| -4 | -92 | 4 | 203 | 0 |
| 40 | -80 | 8 | 134 | 0 |
| -62 | -24 | -14 | 100 | 0 |
| -44 | -46 | -14 | 100 | 0 |
| 8 | -60 | 68 | 67 | 0.000007 |
| 14 | -92 | 0 | 63 | 0.000012 |
| 38 | -52 | 48 | 60 | 0.000018 |
| -30 | -52 | 40 | 57 | 0.000026 |
| 6 | 54 | 16 | 52 | 0.000055 |
| -10 | -52 | 28 | 48 | 0.000101 |
| -12 | -66 | 8 | 45 | 0.00016 |
| -6 | -32 | 28 | 43 | 0.000206 |
| 8 | 8 | 6 | 43 | 0.000206 |
| 20 | 8 | 0 | 41 | 0.00028 |
| 24 | -52 | 6 | 39 | 0.000383 |
| -30 | -2 | 56 | 38 | 0.000426 |
| 46 | -58 | -26 | 38 | 0.000426 |
| 26 | -38 | -16 | 32 | 0.001193 |
| -6 | -18 | 8 | 32 | 0.001193 |
| -24 | -40 | -8 | 27 | 0.003037 |
| -44 | 4 | -36 | 23 | 0.006664 |
| -10 | -66 | -16 | 22 | 0.007996 |
| 56 | -24 | 32 | 21 | 0.009372 |
| 50 | -30 | 56 | 21 | 0.009372 |
| -8 | 58 | 12 | 18 | 0.017283 |
| -6 | 50 | 10 | 18 | 0.017283 |
| 34 | -52 | -14 | 17 | 0.02019 |
| -32 | 16 | -16 | 17 | 0.02019 |
| 8 | -28 | 30 | 17 | 0.02019 |
| -2 | -4 | 20 | 16 | 0.023298 |
| -40 | -48 | 42 | 16 | 0.023298 |
| 6 | -14 | 30 | 16 | 0.023298 |
| 8 | -34 | -28 | 16 | 0.023298 |
| -48 | -36 | 50 | 15 | 0.02896 |
| -32 | -36 | -18 | 14 | 0.034761 |
| 18 | -82 | 12 | 14 | 0.034761 |
| -6 | -70 | -26 | 14 | 0.034761 |

## Discussion

Cognitive models of speech comprehension ^50–52^ have proposed the existence of an auditory lexicon. More recent models of speech comprehension in the brain ^2,3^ have posited the existence of such an auditory lexicon in an auditory ventral “what” stream. Yet, significant disagreement exists between these models about the location of such an auditory lexicon, and no studies to date have directly tested the existence of an auditory lexicon for speech comprehension in the brain. In prior work ^19,20,44^, we have used fMRI rapid adaptation techniques to establish the existence of an orthographic lexicon in a region of the visual ventral stream known as the Visual Word Form Area, VWFA (subsequently confirmed by human neurophysiological recordings ^21,22,53^). In the present study, we leveraged these techniques to test the existence of an auditory lexicon in the auditory ventral stream, particularly in the “Auditory Word Form Area” (AWFA) of the anterior STG ^26,31^. We first defined the ROI for an individual auditory word form area (AWFA) through an independent auditory localizer scan. We then showed in RW and UTPW RA scans that, consistent with an auditory lexicon, spoken RWs engaged distinct neural populations in the AWFA, with even a single phoneme change causing full release from adaptation, whereas pseudowords did not exhibit this lexical adaptation profile but instead showed a graded release from adaptation as a function of phoneme overlap. This graded release from adaptation suggests that neurons in the AWFA, like the VWFA ^20^, exhibit broader tuning to pseudowords due to experience-driven refinement of tuning of neurons to real words but not pseudowords. Thus, these results directly replicated analogous findings of highly selective lexical tuning to real words but not pseudowords in the VWFA ^20^. Then, in the TPW RA scan, we showed that training subjects to recognize a set of pseudowords led to the development of lexical selectivity for these TPWs, again replicating previous results for written words in the VWFA ^19^. Finally, the AWFA was connected to an area in the left posterior fusiform cortex coincident with the reported location of the VWFA, further supporting analogous roles of the AWFA and VWFA in the processing of spoken and written words, respectively.

Our novel evidence of auditory lexical representations in the brain informs current cognitive models of speech comprehension. While all these models map acoustic input to meaning, they disagree on whether auditory lexica exist ^18,54^. Our data do not only present compelling evidence for the existence of an auditory lexicon – they additionally place its location in the anterior STG where it is ideally suited to interface with semantic representations located further anteriorly in the temporal lobe ^16,55^, thereby completing the mapping of speech sounds to meaning. Furthermore, this anterior STG location is consistent with prior studies demonstrating other familiar “auditory objects” ^56^ such as human voices ^57–59^ or musical instruments ^60^. Moreover, such a simple-to-complex progression in selectivity from simple perceptual features over lexical representations to semantic representations in the anteroventral auditory processing stream along the superior temporal cortex is a direct counterpart of the ventral visual stream in the inferior temporal cortex ^9,14^, revealing convergent processing strategies across speech modalities.

## Materials and Methods

### Overall Procedure

In this study, participants completed an auditory localizer scan and three RA experiments over the course of three scanning sessions. In the first session, subjects completed the auditory localizer and real word (RW) scans. The auditory localizer scan was used to identify a candidate AWFA region in the anterior auditory ventral stream^2^ – adopting the approach used in the visual/orthographic case for defining the VWFA ^19,20,44^. The RW RA scan was used to test if the candidate AWFA exhibited lexical selectivity for RWs. In the remaining two scanning sessions subjects completed a pre- and then a post-training PW scans, respectively, separated by 6 behavioral training sessions outside of the scanner.

During the pre-training scan, subjects were presented with the then untrained pseudowords (UTPW). Next, subjects completed six behavioral training sessions consisting of a 2-back and a novel/familiar task designed to familiarize and assess subject familiarity with the UTPW, respectively. Subjects completed a maximum of one behavioral session each day and those participants that achieved at least 80% accuracy on the novel/familiar task by the final session qualified for the post-training scan. During the post-training scan subjects were presented with the same set of pseudowords, now called trained pseudowords (TPW). The pre- and post-training scans were used to test if the candidate AWFA exhibited lexical selectivity for TPW (i.e., after training) but not to the UTPW (i.e., before training).

### Participants

We recruited a total of 28 right-handed healthy native English speakers for this study (ages 18-34, 12 females). This sample size was based on power analyses conducted on prior studies. In two prior studies (Glezer et al., 2015, 2009) that used fMRI rapid adaptation to test the existence of a visual lexicon the effect size (Cohen’s *d*) for the crucial paired t-test that differentiated RWs and UTPWs *d =* 0.87 and 0.69. In another study (Chevillet et al., 2013) that used fMRI rapid adaptation to test phoneme category invariance the effect size (Cohen’s *d*) for the paired t-test between the adaptation for within versus across category exemplars was *d* = 0.93. Using G*Power 3.1 (dependent samples t-test, minimal effect size = 0.69, two-tail α = 0.05, and power = 0.8) found that the sample size is 19. Georgetown University’s Institutional Review Board approved all experimental procedures, and written informed consent was obtained from all subjects before the experiment. Two subjects were excluded from further analyses for performing 2 standard deviations below the average on the in-scanner task. Furthermore, two subjects dropped out of the study after completing the RW RA scan. In total, we analyzed 26 subjects for the RW RA scan and 24 of those 26 for the pre-training UTPW scan. Due to subject dropout (n=4) or failure to achieve 80% accuracy on the novel/familiar task (n=4), 16 out of the 24 subjects were analyzed for the post-training TPW scan.

### Stimuli

Real word stimuli, for the RW RA experiment, were chosen using the English Lexicon Project ^61^. Analogous to our studies of the neural representation of written words ^19,20^, three sets of fifty high-frequency (>50 per million), monosyllabic RWs that were 3-5 phonemes in length were created. One set of words (target words) served as the reference for the other two lists. The second set was created by altering each target word by a single phoneme to create another real word. The third set was created by selecting for each target word another real word that had the same number of (but no shared) phonemes. All three of these lists were matched on the number of phonemes, orthographic neighborhood, and phonological neighborhood. To create the UTPW/TPW we used MCWord (Medler, D.A., & Binder, 2005) to generate four sets of 50 target PWs, 3-5 phonemes in length. One set to be trained and three sets to remain untrained and serve as foils in the training task. As with the RW stimuli, we then used the target PW as the reference to generate a set of pseudowords each differing by one phoneme from the target word, and another set of pseudowords matched to the target pseudowords by number of phonemes but not sharing any phonemes. As in prior work ^19,20^ real word and pseudoword sets were matched for length, bigram and trigram frequency, and phonological neighborhood. All stimuli were recorded using a 44.1-kHz sampling rate in a sound-isolated booth by a trained female native speaker of American English.

### Auditory Localizer Scan

The auditory localizer scan was used to independently identify auditory speech-selective areas. In this scan, subjects were randomly presented with trials from one of five conditions: “Real Words”, “Pseudowords”, “Scrambled Real Words”, “Scrambled Pseudowords”, and “Silence”. “Real Words” and “Pseudowords” were one syllable long; the lists were matched for length, orthographic and phonologic neighborhood. “Scrambled Real Words” and “Scrambled Pseudowords” were generated by randomly rearranging 200 ms by 1-octave tiles of the constant Q spectrogram^62^ for each real word or pseudoword, respectively, and then reconstructing a time-domain waveform with an inverse transform^33^. Each trial in the auditory localizer scan was 9s long and began with 2.5 s of silence (during which 1.68 s of scanner acquisition occurred) followed by a ∼3-4 s stimulus presentation period and concluding with silence. During the stimulus presentation period, subjects heard four stimuli from a given condition and responded with a left-handed button press if any of the four stimuli was a repeat within that block. In total, there were 145 trials per run with 25 trials of each of the five conditions and five one-back trials for each of the four non-silence conditions. An additional 18 s of fixation were added to the start and end of each run. Subjects completed five runs of the task.

### Rapid Adaptation (RA) Scans

There were three RA scans (i.e., RW, UTPW, TPW) performed on different days. In the rapid adaptation (RA) scans, subjects heard a pair of words (prime/target) on each trial. The words in each pair were either identical (SAME), differed by a single phoneme (1PH), or shared no phonemes at all (DIFF). These pairs were generated by using the three matched word lists described above. To engage subjects’ attention, we asked subjects to perform an oddball rime detection task in the scanner. To do so, we created an additional condition (oddball) in which a word or pseudoword containing the oddball rime ‘-ox’ (e.g., socks, grox) was presented in lieu of either the prime or target word. Participants were asked to attentively listen to all stimuli and respond with a left-handed button press when an oddball stimulus was heard. In all three scans, the number of repetitions of each word was counterbalanced across all conditions to control for long-lag priming effects^63^. Trial order and timing was adjusted using M-sequences^64^. Each trial was 3.36 s long and consisted of a 1.68 s silent period followed by 1.68 s stimulus presentation period during which the word pairs were presented. Following prior auditory RA studies^45,46^ we presented stimuli with a 50ms ISI. In total, there were 25 trials of each condition (SAME, 1PH, DIFF, oddball, and silence) for a total of 125 trials per run. An additional 10.08 s of fixation was added to the start and end of each run. Subjects completed 4 runs for each scan.

### Data Acquisition

MRI data were acquired at the Center for Functional and Molecular Imaging at Georgetown University on a 3.0 Tesla Siemens Prisma-fit scanner. We used whole-head echo-planar imaging sequences (flip angle = 70°, TE = 35 ms, FOV = 205mm, 102×102 matrix) with a 64-channel head coil. Building off previous auditory localizer^65^ and RA paradigms in our lab ^46^, a slow (TR = 9000 ms, TA = 1680 ms) and a fast (TR = 3360 ms, TA = 1680 ms) clustered acquisition paradigm were used for the auditory localizer and RA scans, respectively. 54 axial slices were acquired in descending order (thickness = 1.8 mm, 0.18 mm gap; in-plane resolution = 2.0 × 2.0 mm^2^). A T1-weighted MPRAGE image (resolution 1×1×1 mm^3^) was also acquired for each subject.

### fMRI Data Preprocessing

Image preprocessing was performed using SPM12 (http://www.fil.ion.ucl.ac.uk/spm/software/spm12/). The first three volumes of each run were discarded to allow for T1 stabilization, and the remaining EPI images were spatially realigned to the mean BOLD reference image. No slice-time correction was done given the presence of temporal discontinuities between successive volumes in clustered acquisition paradigms ^66^. EPI images for each subject were co-registered to the anatomical image. The anatomical image was then segmented and the resulting deformation fields for spatial normalization were used to normalize the functional data to the standard MNI space. Next, we smoothed the normalized functional images with a 4-mm FWHM Gaussian kernel. Finally, a first-level model containing regressors for each condition and the six motion parameters from realignment was fit. In the auditory localizer scan, the regressors were: “Real Words”, “Pseudowords”, “Scrambled Real Words”, “Scrambled Pseudowords”, “Button Press”, and “Silence”. In the RA scans, the regressors were: “SAME”, “1PH”, “DIFF”, “Button Press”, and “Silence”.

### Defining the Auditory Word Form Area (AWFA)

The auditory word form area (AWFA) was determined at the individual subject-level using the auditory localizer scan. Analogous to our studies of the visual word form area ^19,20,44^, for each subject we first defined an “*RW-Silence”* contrast and an “*RW-Scrambled Words”* contrast. Next, at the group level we masked the “*RW-Silence”* t-statistic map with the “*RW-Scrambled Words”* t-statistic map thresholded at p ≤ 0.05 before applying a p ≤ 0.001 voxel-wise threshold. The resulting map had a peak at MNI: -62, -14, 2. These coordinates are consistent with the hypothesized locus (MNI: -61, - 15, -5) of the AWFA from prior studies ^26^. We then created the same “*RW-Silence”* masked by “*RW-Scrambled Words” maps at the* individual-subject level using the same thresholds as above. Then, for each subject we identified the local peak in the resulting maps closest to the group peak (MNI: -62, -14, 2). Finally, to create each individual subject’s AWFA, we created a region of interest (ROI) consisting of the 50 closest voxels to each individual’s local peak.

### Behavioral Training

Subjects were trained to recognize 150 auditory PWs (TPW, see above). Following previous work^19^, a 2-back training task, in which subjects had to detect repeats of PW separated by another PW, was used to familiarize subjects with the PWs. Each session of the 2-back task consisted of 15 blocks of 75 trials each with self-paced breaks between each block. Each trial lasted 1.5 s during which subjects heard a single PW and had to respond if they heard a 2-back repeat (i.e., if the current PW was the same as the PW before the last). Each block lasted 112.5 s, and each session lasted for a total task length of 28.125 min excluding breaks. Following the 2-back task, subjects’ familiarity with the trained PWs was assessed using a novel/familiar task. For this task, we developed 3 sets of foils for each of the 150 PWs. Each foil differed from its base PW by a single phoneme. Each session of the novel/familiar task consisted of 3 blocks of 100 trials with self-paced breaks between each block. Each trial lasted 1.5 s during which subjects heard a single PW and had to respond with either the ‘left’ or ‘right’ arrow key to indicate a novel (i.e., a foil) or familiar (i.e., trained) PW. In total, each block lasted 150 s for a total task length of 7.5 min excluding breaks. Over the course of the novel/familiar task sessions each foil list was paired with the trained PW list only twice with at least 2 days since the last pairing. Six sessions of both tasks were performed over the course of ∼8 days. To proceed to the post-training rapid adaptation (RA) scan, subjects had to achieve at least 80% accuracy on the novel/familiar task by their 6^th^ session.

### Task-Based Functional Connectivity

After preprocessing, we used the CONN toolbox 21.a ^67^ to calculate seed-to-voxel stimulus-driven functional connectivity from the AWFA during auditory word processing in the auditory localizer scans. Word processing was included as the primary task condition by combining the onset times and durations for RW and PW into one condition. The individual subject (n=26) AWFA seeds were used for the analysis. We then performed denoising with confounds including the subject-specific motion regressors computed during preprocessing in SPM, as well as modeling nuisance covariates including cerebrospinal fluid and white-matter signals and their derivatives, following the CompCor strategy ^68^ as implemented in CONN. Data were then band-pass filtered (0.008-0.009 Hz) with linear detrending. Seed-to-voxel functional connectivity was performed using a weighted-GLM computing the bivariate correlation between the AWFA seed with the whole brain. Due to the sparse acquisition type of this analysis, no HRF weighting was performed. Group-level significance was determined with a voxel threshold of p<0.001 and a cluster threshold of p<0.05, FDR-corrected.

## Acknowledgements

We thank Iain DeWitt for providing the auditory scrambling code used for the localizer scans. The research described in this paper was funded by NSF (grant BCS-1756313). Additional support was provided by the NIH (grant 1S10OD023561). This work also used resources provided through the Extreme Science and Engineering Discovery Environment (XSEDE), which is supported by National Science Foundation grant number ACI-1548562

